# *Tet* Trim-Away: A conditional rapid protein degradation system for *Tetrahymena thermophila*

**DOI:** 10.64898/2026.05.29.728803

**Authors:** Gayatri L. Dholakia, Alexander J. Stemm-Wolf, Aaisha Anzar, Aaron P. Turkewitz, Chad G. Pearson

## Abstract

*Tetrahymena thermophila* is a ciliated protist that has played pivotal roles in biological discovery. Functional studies of *Tetrahymena* proteins have largely relied on gene knockouts. Because protein depletion upon knockout typically spans multiple cell cycles, compensatory mechanisms can confound phenotypic interpretation. To instead enable rapid and acute protein depletion, we modified and adapted the Trim-Away system for use in *Tetrahymena (Tet* Trim-Away). Trim-Away is based on the E3 ubiquitin ligase, TRIM21, that binds to antibody-bound proteins and targets them for proteasome mediated degradation. Here, Trim-Away was modified with a fusion of the N-terminal RBCC (RING, B-box, coiled-coil) domains of TRIM21 with an α-mCherry (mCh) nanobody sequence that recognizes endogenously tagged mCh proteins of interest (Nb^mCh^). Expression of the RBCC:Nb^mCh^ degron, which is controlled by an inducible promotor, promotes rapid target protein depletion within 30 minutes and can be sustained for weeks. *Tet* Trim-Away is reversible, functions against targets in multiple cellular compartments, and produces loss-of-function phenotypes in *Tetrahymena* cells.

## INTRODUCTION

*Tetrahymena thermophila* is a eukaryotic ciliate amenable to forward and reverse genetic approaches (Ruehle et al., 2016). Genetic alterations combined with sophisticated biochemical and cell biology studies have made this model organism a driver of discovery for fundamental biological processes such as telomere synthesis, histone modifications, and dynein-based transport. Forward genetic studies have generated mutants leading to new understanding of drug resistance, secretion, cilia, cortical stability, cell patterning, and the cell cycle (Cole and Gaertig, 2022; Cole et al., 2026; Frankel, 2008; Orias et al., 1999; Pennock, 1999; Roberts and Orias, 1973; Stemm-Wolf et al., 2026). Reverse genetic studies have capitalized on the ability to produce gene knockouts via homologous recombination (Ruehle et al., 2016) or by methods that exploit specialized genome editing in ciliates (Hayashi and Mochizuki, 2015). These established methods to study the function of proteins of interest in *T. thermophila* rely on protein depletion that occurs over multiple cell cycles. Importantly, during the extended period of protein depletion, cells can potentially activate compensatory mechanisms that interfere with researchers’ ability to study the function of the protein of interest (Rossi et al., 2015). This limitation highlights the need to develop methods for rapid depletion to more precisely analyze the functions of *Tetrahymena* proteins.

Degron systems have provided effective means for studying protein function. Proteolysis targeting chimeras (PROTACs) enable targeting of specific proteins for depletion using the cell’s proteasomal degradation machinery (Sakamoto et al., 2001). A variation on the PROTAC approach is Trim-Away. Trim-Away uses the tripartite motif-containing protein 21 (TRIM21), an E3 ubiquitin ligase with RING, B-box, and coiled-coil domains (RBCC) at the N-terminus and a PRY/SPRY domain at the C-terminus. The N-terminal RBCC promotes protein ubiquitinylation and degradation. The C-terminal PRY/SPRY domain binds the Fc region of antibodies, thereby targeting proteins recognized by antibodies that are added to the cell (Clift et al., 2017; James et al., 2007; Mallery et al., 2010). A limitation of this system is that it requires a large amount of antibody to the protein of interest. To overcome the need for exogenously supplied antibodies, the PRY/SPRY domain was replaced with nanobody gene fusions (Chen et al., 2021; Fletcher et al., 2023). Nanobodies are small, stable, single-domain fragments derived from the variable heavy chain domain of camelid antibodies, which lack a light chain (He et al., 2025; Ji et al., 2022; Jiang et al., 2024; Prole and Taylor, 2019; Zhang et al., 2021). To create the Trim-Away variant, nanobodies were recombinantly fused to the TRIM21 RBCC motif to generate fusion proteins known as TRIMbodies or TRIM21-based bioPROTACs. When expressed in cells, these recognize the ligand of the nanobody and trigger its proteasome-mediated degradation (Chen et al., 2021; Fletcher et al., 2023). Thus, Trim-Away represents an efficient approach for rapid degradation of selected proteins.

Here, we developed Trim-Away for use in *Tetrahymena* (*Tet* Trim-Away). A *Tetrahymena* codon-optimized N-terminal RBCC motif was fused to an α-mCherry (mCh) nanobody (RBCC:Nb^mCh^), and this was assembled within a construct that allows for inducible expression in *Tetrahymena.* We introduced this construct into cells co-expressing a variety of genes that were each endogenously tagged with mCh. Induction of RBCC:Nb^mCh^ rapidly and specifically degraded mCh-tagged proteins within 30 minutes with sustained degradation persisting for weeks. We show that *Tet* Trim-Away depleted proteins in distinct cellular compartments, degrading cytoplasmic, ciliary, and nuclear proteins. Moreover, degradation was reversible and produced strong depletion phenotypes. Therefore, *Tet* Trim-away offers a selective degron system that should facilitate more precise functional studies of *Tetrahymena* proteins.

## RESULTS

### *Tet* Trim-Away depletes Poc1:mCh protein

The N-terminal RBCC motif of *H. sapiens* TRIM21 was codon optimized for *Tetrahymena* and fused with a nanobody specific to mCh (RBCC:Nb^mCh^) (Supplemental Figure 1A; (Chan et al., 1991; Fridy et al., 2014; Holzer et al., 2022; Itoh et al., 1991; Tsugu et al., 1994); Protein Accession NP_003132). This was subcloned into a construct whose expression in *Tetrahymena* is controlled by CdCl_2_ induction of the metallothionein (*MTT1*) promoter (Piccinni et al., 1999; Shang et al., 2002). To test whether expression of RBCC:Nb^mCh^ results in degradation of mCh-tagged proteins, the construct was transfected into cells expressing endogenously tagged Poc1:mCh (Pearson et al., 2009). Poc1 protein localizes to basal bodies, the sites of ciliary axoneme organization. RBCC:Nb^mCh^ was induced with 5.5 μM (1.0 μg/mL) CdCl_2_ in SPP media. This concentration of CdCl_2_ induced strong depletion of Poc1:mCh without cellular toxicity (Figure 1 and Supplemental Figure 1B-D). 30 minutes post RBCC:Nb^mCh^ induction, 68±5% of cells exhibited nearly undetectable Poc1:mCh signal by fluorescence microscopy (Figure 1A, B; Low Signal). The mean fluorescence signal was 30±9% of the control (Figure 1C). 45 minutes after RBCC:Nb^mCh^ induction, 99±2% cells exhibited low fluorescence signal and the mean fluorescence intensity was reduced to 16±4% (Figure 1B, C). Three hours after RBCC:Nb^mCh^ induction, 23±33% of the cells had no detectable Poc1:mCh signal. The mean fluorescence signal was reduced to 14±6% (Figure 1A, D, E). This combination of low or no signal observed in the cells persisted through 336 hours of induction (Figure 1A, D, E and Supplemental Figure 2A and B). Thus, *Tet* Trim-Away produces rapid onset protein degradation that is also persistent.

**Figure 1.**
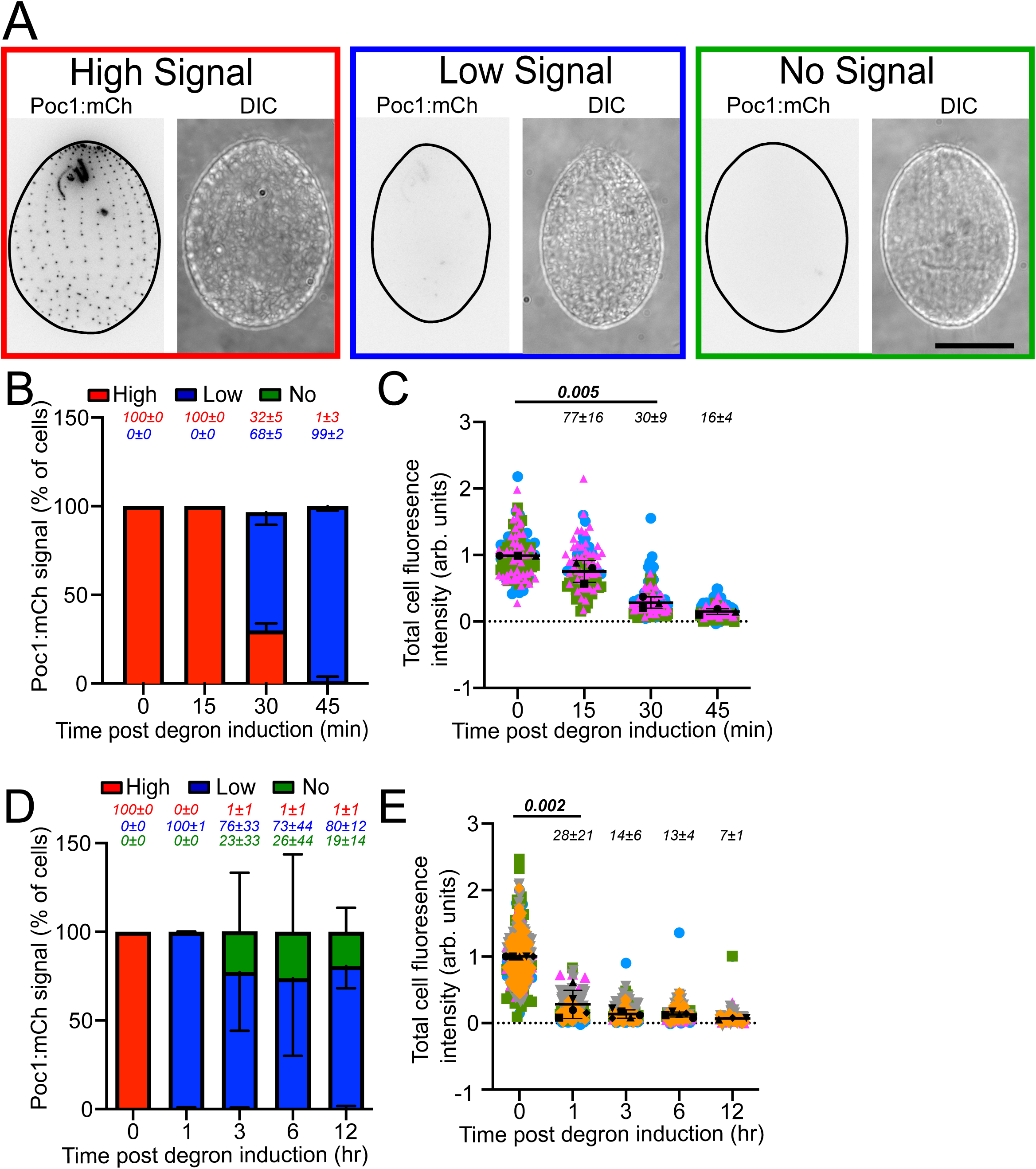
*Tet* Trim-Away for Poc1:mCh. (A) Images of Poc1:mCh cells expressing RBCC:Nb^mCh^. Cells were classified into high (outlined in red), low (outlined in blue), and no detectable Poc1:mCh signal (outlined in green) (left, mCh fluorescence; right, differential inference contrast (DIC)). (B) Qualitative analysis of RBCC:Nb^mCh^ Trim-Away Poc1:mCh degradation through 45 minutes of induction. Cells were classified as described in (A) and percentages for each class are shown in the graph. (C) Quantitative analysis of RBCC:Nb^mCh^ Trim-Away degradation of Poc1:mCh. Whole cell Poc1:mCh fluorescence intensity was measured through 45 minutes of induction. Each data point represents a cell and mean fluorescence intensity of the time point is indicated above the data points (as a percentage). (D) Qualitative analysis of RBCC:Nb^mCh^ Trim-Away of Poc1:mCh degradation through 12 hours of induction. Percentages for each class are shown in the graph. (E) Quantitation of whole cell Poc1:mCh fluorescence intensity through the 12 hour time course of RBCC:Nb^mCh^ mediated degradation. Percentage of mean fluorescence intensity relative to control of each time point is indicated above the data points. Error bars indicate mean±SD. Scale Bars: 20μm (B) *N > 300* for qualitative analysis and (C) *N > 120* for quantitative analysis (D) *N > 500* for qualitative analysis and (E) *N> 200* for quantitative analysis.

In a related construct, we replaced the nanobody in our *Tet* Trim-Away construct with one specific for GFP rather than mCh, to create RBCC:Nb^GFP^. We tested its ability to induce degradation of Poc1:GFP, using the same approach as that described for Poc1:mCh expressing cells. Three hours post RBCC:Nb^GFP^ induction, 50±6% of cells maintained high Poc1:GFP signal, 48±6% of cells exhibited low signal and 2±2% cells had no detectable signal. These ratios were maintained through 12 hours of degron induction. The mean fluorescence signal was reduced to 59±22% of control cells throughout the time course. Compared to the response induced by RBCC:Nb^mCh^ *Tet* Trim-Away, the degradation induced by RBCC:Nb^GFP^ was less effective (Supplemental Figure 2C and D). In summary, our results in the context of Poc1 fusion proteins indicate that of the two *Tet* Trim-Away constructs we have tested, RBCC:Nb^mCh^ *Tet* Trim-Away appears highly effective while RBCC:Nb^GFP^ requires further optimization.

### *Tet* Trim-Away mediated depletion is reversible

To determine whether rapid targeted depletion of Poc1:mCh is reversible, Poc1:mCh was first depleted by a six-hour induction of RBCC:Nb^mCh^, resulting in 73±23% cells with low signal and 27±23% cells with no detectable signal, and 29±13% mean fluorescence intensity compared to control cells (Figure 2). The cells were then washed of CdCl_2_ and suspended in fresh medium. By three hours after washout, high Poc1:mCh signal was detectable in 15±25% of cells, 77±21% cells exhibited low and 8±11% cells had no detectable signal (Figure 2A and B). By 12 hours, 96±7% cells exhibited high Poc1:mCh signal. The mean fluorescence signal intensity increased from 24±21% at three hours to 88±32% by 12 hours. These results demonstrate that washout of CdCl_2_ suffices to restore accumulation of Poc1, so depletion is reversible. Interestingly, at 24 hours, the cells overcompensated with a mean fluorescence signal elevated to 162±47% of the original levels (Figure 2A and B).

**Figure 2.**
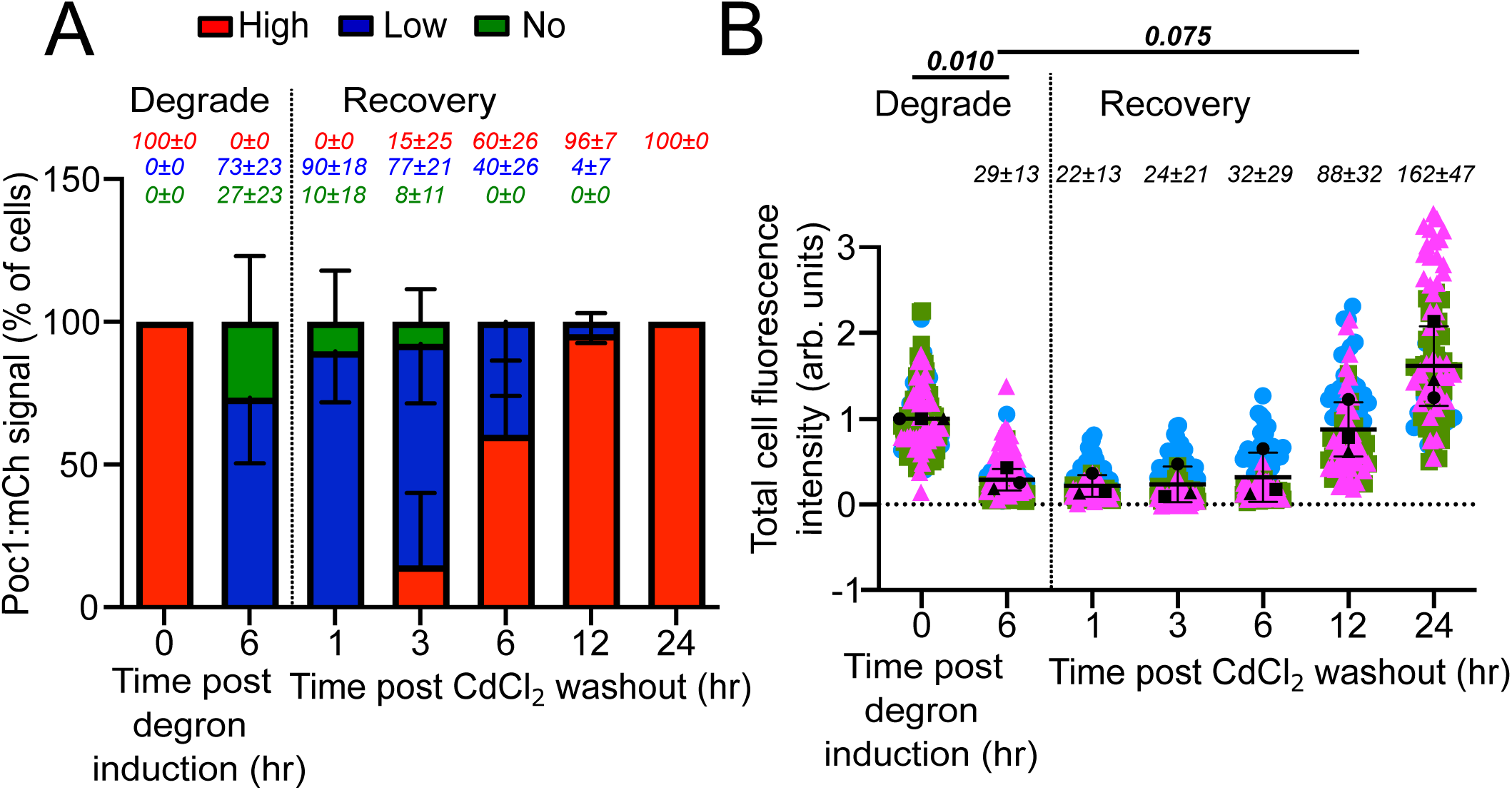
*Tet* Trim-Away is reversible. (A) Qualitative analysis of recovery after washout of RBCC:Nb^mCh^ mediated Poc1:mCh degradation. RBCC:Nb^mCh^ Trim-Away was expressed for six hours to degrade Poc1:mCh. CdCl_2_ was washed from the media and cells were fixed at one, three, six, 12 and 24 hours after washout. (B) Whole cell Poc1:mCh mean fluorescence intensity through the washout experiment. Percentages of each class and mean fluorescence intensity of each time point relative to control are indicated above the data points. Error bars indicate mean±SD. *N > 300* for qualitative analysis and *N > 120* for quantitative analysis.

### *Tet* Trim-Away requires proteasome activity and localizes with the target protein

To test whether proteasomal activity is required for induced Poc1:mCh degradation, we took advantage of the proteasome inhibitor MG132. Cells were treated with 50-150 μM MG132 for 45 minutes and subsequently exposed to CdCl_2_ for one hour to induce RBCC:Nb^mCh^ expression. 47±6% of cells treated with 50 μM MG132 retained high Poc1:mCh signal, indicating that inhibition of proteasome activity resulted in fewer cells degrading Poc1:mCh. Mean fluorescence intensity was 42±25% of control cells as compared to 12±3% for cells without MG132 treatment (Figure 3A and B). The inhibition of depletion of Poc1:mCh by MG132 was dose dependent. These results indicate that *Tet* Trim-Away-mediated depletion depends upon proteasome degradation activity. Furthermore, 150 μM MG132 increased the mean fluorescence of Poc1:mCh to 159±26% of control levels. The elevation of Poc1:mCh beyond control levels, and that MG132 alone does not increase basal levels of Poc1:mCh (Supplemental Figure 3), suggests a compensating upregulation of Poc1:mCh expression in response to *Tet* Trim-Away.

**Figure 3.**
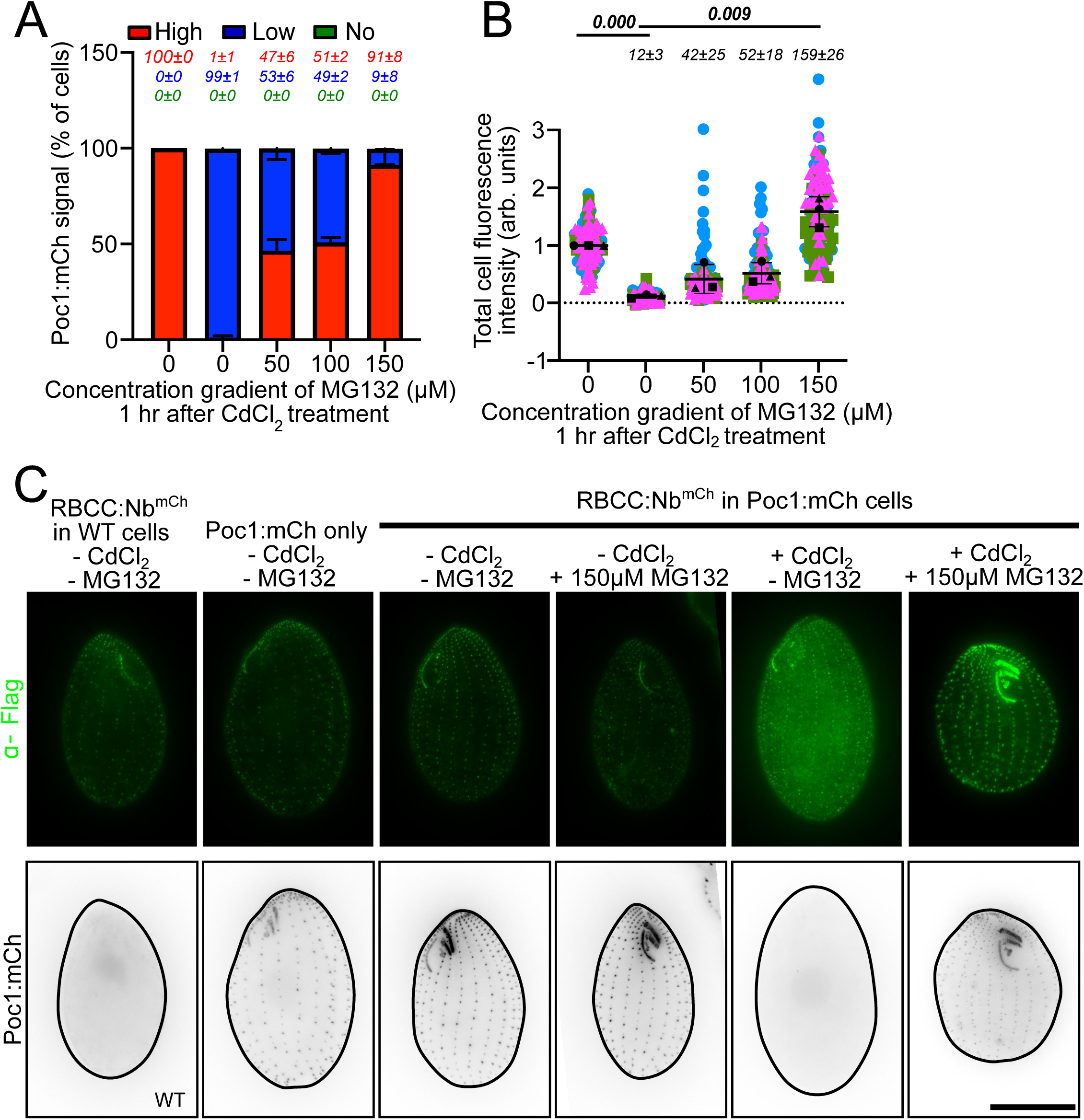
*Tet* Trim-Away requires proteasome activity and localizes to basal bodies. (A) Qualitative analysis of inhibition of RBCC:Nb^mCh^ mediated Poc1:mCh degradation by blocking the proteasome using MG132. (B) Whole cell Poc1:mCh mean fluorescence intensity with different concentrations of the proteosome inhibitor, MG132. Percentages of each class and mean fluorescence intensity, expressed as percentages of each time point relative to control are indicated above the data points. (C) α-Flag immunofluorescence staining (green) for wildtype cells, Poc1:mCh cells without RBCC:NB^mCh^, and cells with Poc1:mCh and RBCC:Nb^mCh^. Poc1:mCh signal is shown in grey scale. MG132 was added to cells 45 minutes before addition of CdCl_2_. Cells were fixed and imaged one hour after CdCl_2_ treatment. Scale bar, 20μm. Error bars indicate mean±SD. (A, B) *N > 300* for qualitative analysis and *N > 120* for quantitative analysis.

To establish where in the cell the RBCC:Nb^mCh^ binds Poc1:mCh, RBCC:Nb^mCh^ was localized using immunofluorescence staining to the 1X Flag tag present on RBCC:Nb^mCh^ (Supplemental Figure 1A). In wildtype and Poc1:mCh cells not expressing RBCC:Nb^mCh^, dim α-Flag immunofluorescence signal was observed at basal bodies (Figure 3C and Supplemental Figure 3A-C). Therefore, a low level of α-Flag antibody localizes non-specifically to basal bodies. Upon induction of RBCC:Nb^mCh^ in Poc1:mCh cells, α-Flag fluorescence signals grew brighter throughout the cell, including at basal bodies. The fluorescence at basal bodies in these cells further intensified upon addition of 150 μM MG132 (Figure 3C). The localization of RBCC:Nb^mCh^ in Poc1:mCh cells to basal bodies suggests that RBCC:Nb^mCh^ is effectively recruited to its target protein sites.

### *Tet* Trim-Away targeting is highly specific

To test whether RBCC:Nb^mCh^ *Tet* Trim-Away expressed in Poc1:mCh cells promotes non-specific targeting and degradation of other proteins within the basal body, centrin that resides in basal bodies was visualized using α-centrin staining (Stemm-Wolf et al., 2005). One and six hours post *Tet* Trim-Away induction, Poc1:mCh signal was significantly decreased but no change in centrin signal intensity or localization was observed (Figure 4A-C). Similarly, Poc1:mCh degradation did not affect associated basal body tubulin as measured by α-acetylated tubulin staining (Figure 4D-G). Importantly, in these experiments we did not provoke the global basal body defects associated with Poc1 loss because in these cells the tagged *POC1:mCH* allele only partially replaced the endogenous gene and therefore untagged alleles of *POC1* could support normal basal body structure and stability. To conclude, *Tet* Trim-Away tailored for Poc1:mCh does not target nearby basal body proteins and does not promiscuously disrupt the basal body structure.

**Figure 4.**
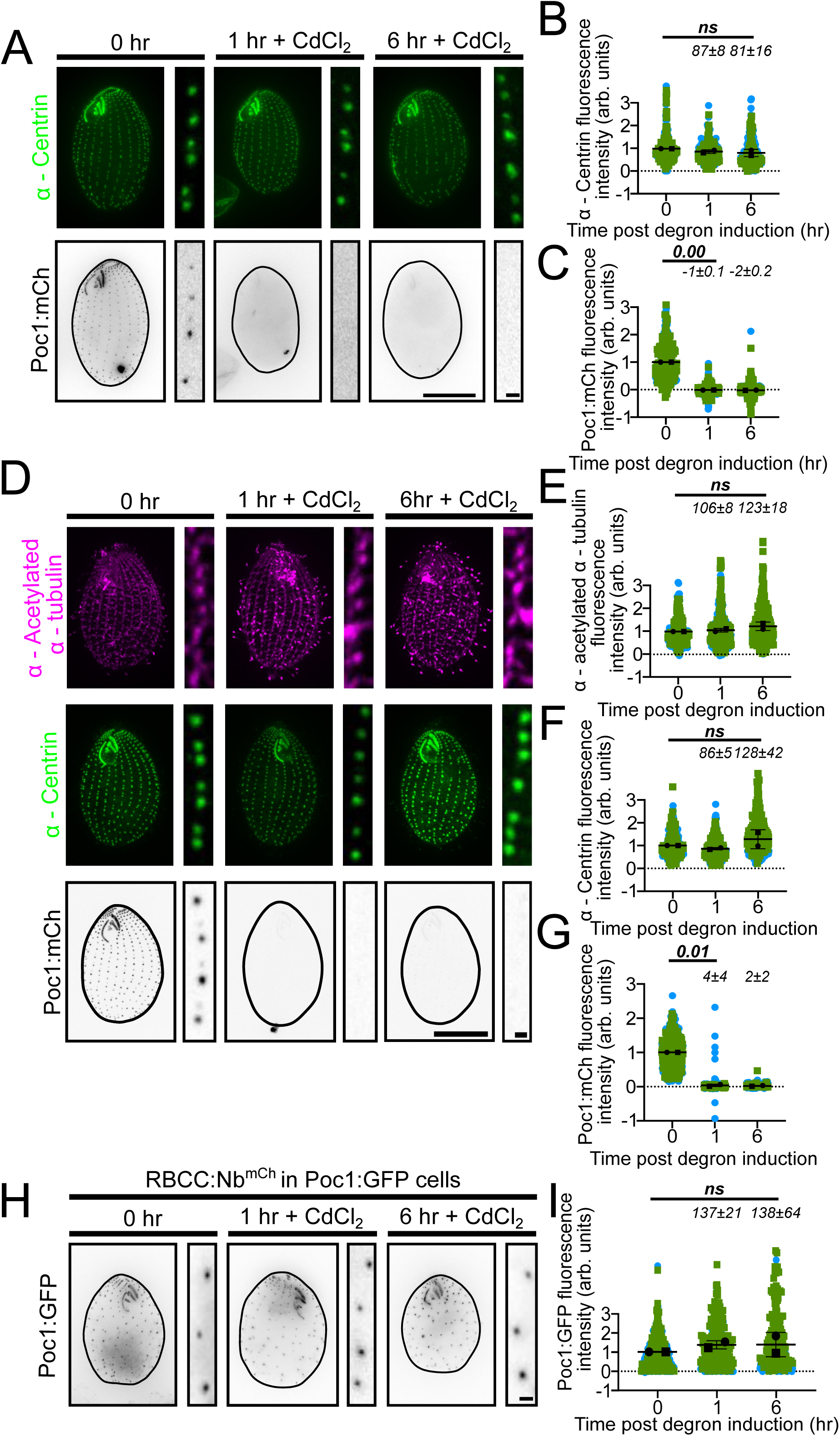
*Tet* Trim-Away specifically degrades Poc1:mCh. (A) α-Centrin localized to basal bodies (green). Poc1:mCh signal shown in grey scale. (B) Fluorescence intensity of centrin signal at basal bodies through six hours of RBCC:Nb^mCh^ induction. (C) Poc1:mCh fluorescence intensity at basal bodies. (D) α-acetylated α-tubulin staining shown in magenta, α-Centrin localization to basal bodies in green, and Poc1:mCh in grey scale. (E) α-acetylated α-tubulin fluorescence intensity signal at basal bodies. (F) Fluorescence intensity of α-centrin signal at basal bodies. (G) Poc1:mCh fluorescence intensity at basal bodies through six hours of RBCC:Nb^mCh^ expression. (H) RBCC:Nb^mCh^ expression in Poc1:GFP cells. Poc1:GFP signal in grey scale. (I) Fluorescence intensity of Poc1:GFP at basal bodies of cells through six hours of RBCC:Nb^mCh^ induction. Mean fluorescence intensity of each time point relative to control are indicated above the data points (as percentages). Scale bars, 20μm; 1μm. Error bars indicate mean±SD. (B,C,E,F,G,I) *N > 300* for quantitative analysis.

As a further test of the specificity of the α-mCh nanobody, RBCC:Nb^mCh^ was induced in cells expressing Poc1:GFP. After 6 hours of RBCC:Nb^mCh^ expression, Poc1:GFP signal was undiminished, and the mean fluorescence intensity of basal bodies was not significantly changed (Figure 4H and I). This suggests that RBCC:Nb^mCh^ is specific to the mCh fusion protein and does not target GFP-tagged proteins.

### *Tet* Trim-Away effectively degrades proteins in diverse cellular compartments

To be maximally useful, a tool for induced protein degradation should function in a wide range of cellular compartments. To test the efficacy of RBCC:Nb^mCh^ *Tet* Trim-Away in the context of cellular structures and compartments besides the ciliary basal bodies, we engineered cells expressing a variety of mCh-tagged fusion proteins. In particular, we expressed RBCC:Nb^mCh^ in cells that co-expressed an mCh-tagged cortical striated fiber protein (Bbc39), ciliary axoneme radial spoke protein (Rsph9), or H3 nuclear histone (Hht2).

Bbc39 localizes to striated fibers in the cortical cytoskeleton, which attach to basal bodies. Such proteins are required for building and maintaining *Tetrahymena* cortical architecture (Frankel, 1999; Frankel, 2008; Galati et al., 2014; Soh et al., 2020). Within one hour of RBCC:Nb^mCh^ expression, Bbc39:mCh cells exhibited a 98±1% loss of mCh signal. The mean fluorescence intensity diminished to an average of 9±2% that of control cells, and that dramatic depletion persisted through 12 hours of RBCC:Nb^mCh^ expression (Figure 5A-C). When whole cell Bbc39:mCh degradation was measured by immunoblotting of cell lysates, the protein levels were found to be reduced to 5±2% and 6±7% at one and six hours after RBCC:Nb^mCh^ induction, respectively (Supplemental Figure 4A and B). In summary, *Tet* Trim-Away effectively depletes a striated fiber protein. Interestingly, the low-level residual signal persists at elongated fibers that are displaced from basal bodies and elongate with time, suggesting this protein population becomes resistant to degradation (Figure 5A).

**Figure 5.**
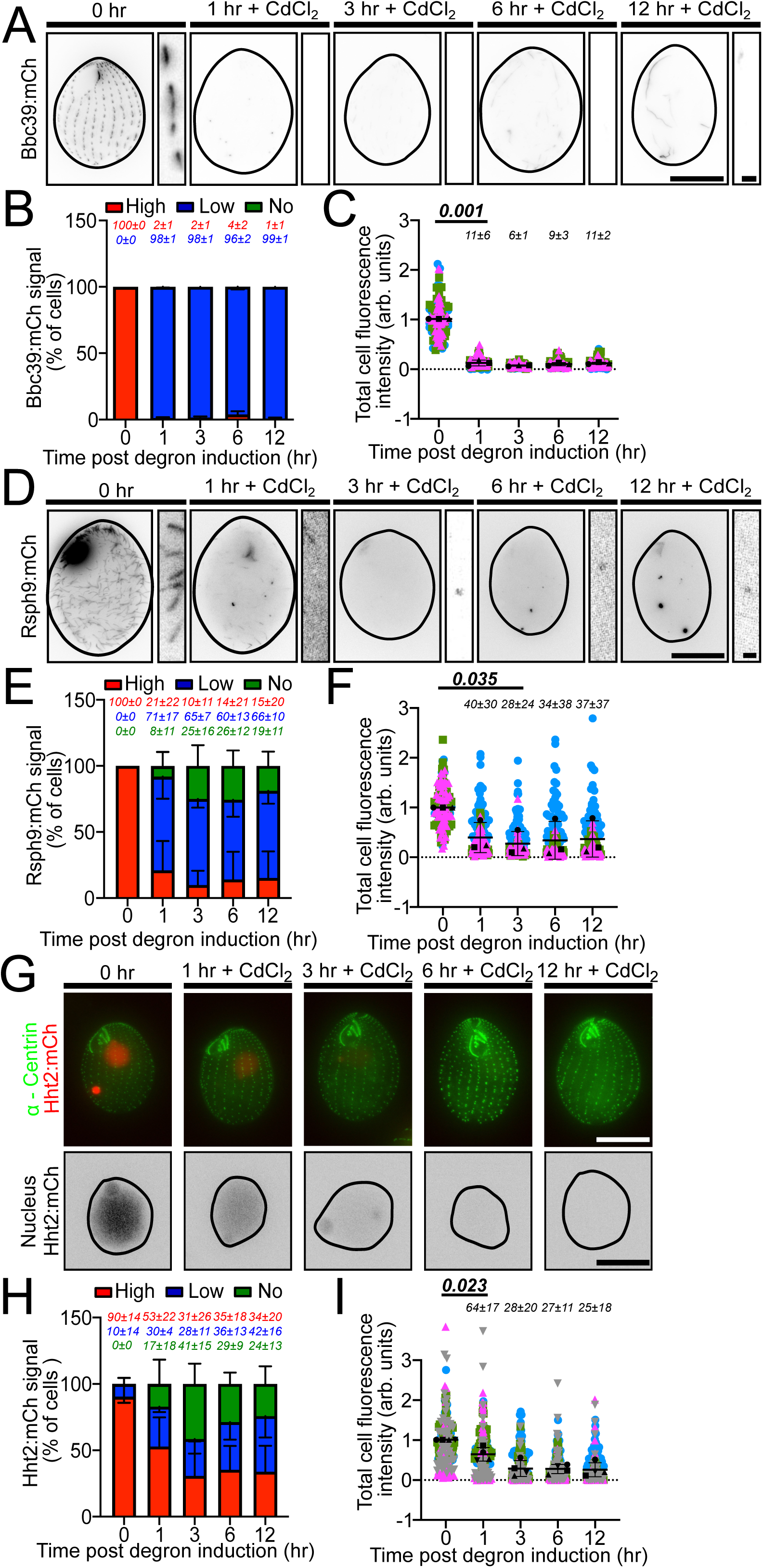
*Tet* Trim-Away degrades proteins in diverse cellular compartments. (A) Bbc39:mCh expressing RBCC:Nb^mCh^ at 0, one, three, six, and 12 hours post RBCC:Nb^mCh^ Trim-Away induction. Bbc39:mCh in grey scale. (B) Qualitative analysis of Bbc39:mCh degradation. (C) Whole cell Bbc39:mCh mean fluorescence intensity through 12 hours of RBCC:Nb^mCh^ Trim-Away expression. (D) Rsph9:mCh (grey scale) cells expressing RBCC:Nb^mCh^. Image of cell at 0 hour is brighter than rest of the images. The image was not saturated when captured. (E) Qualitative analysis of Rsph9:mCh degradation through 12 hours of RBCC:Nb^mCh^ induction. (F) Rsph9:mCh whole cell mean fluorescence intensity through 12 hours of RBCC:Nb^mCh^ Trim-Away expression. Scale bars, 22.5 μm; 1μm. (G) Hht2:mCh cells expressing RBCC:Nb^mCh^. Cells stained with α-Centrin (green) and Hht2:mCh (red) (top). Macro and micronuclei labelled with Hht2:mCh in grey scale (bottom). (H) Qualitative analysis of Hht2:mCh degradation through 12 hours of RBCC:Nb^mCh^ induction. (I) Hht2:mCh nuclear mean fluorescence intensity through 12 hours of RBCC:Nb^mCh^ Trim-Away expression. Percentages of each depletion class and mean fluorescence intensity of each time point relative to control are indicated above the data points. Scale bar, 20μm, 5μm. Error bars indicate mean±SD. (B, C, E, F) *N > 300* for qualitative analysis and *N > 120* for quantitative analysis. (H, I) *N > 400* for qualitative analysis and *N > 160* for quantitative analysis.

Radial spoke-head protein 9 (Rsph9) is a ciliary component that promotes normal ciliary beating (Castleman et al., 2009; Narita et al., 2012; Yang et al., 2006). A three hour induction of RBCC:Nb^mCh^ expression in Rsph9:mCh cells resulted in 65±7% cells with low signal and 25±16% cells with no detectable signal (Figure 5D-F). This ratio of cells with low and no signal persisted through 12 hours of RBCC:Nb^mCh^ induction. 15±5% of cells consistently expressed high Rsph9:mCh signal throughout the time course. Cells at six hours of degron expression (mean fluorescence intensity 34±38% of control cells) and 12 hours (mean fluorescence intensity 37±37%) showed some recovery of fluorescent signal compared to cells at three hours (mean fluorescence intensity 28±24%) (Figure 5E and F). In summary, *Tetrahymena* Trim-Away is competent to deplete the Rsph9 ciliary protein, though less completely than the other protein targets tested above.

To test whether Tet Trim-Away can degrade nuclear proteins, RBCC:Nb^mCh^ was expressed in cells with histone H3 fused to mCh (Hht2:mCh). Hht2 localizes to both nuclei present in each *Tetrahymena* cell, the somatic macronucleus (MAC) and the germline micronucleus (MIC). Within one hour of RBCC:Nb^mCh^ expression, 30±4% of cells expressed low signal and 17±18% of cells had no detectable signal (Figure 5G-I). However, 53±22% of cells retained high signal. The mean fluorescence intensity was reduced to 64±17% of control cells. By six hours, 36±13% cells had low signal and 29±9% cells had no detectable signal (mean fluorescence intensity reduced to 27±11%). This ratio persisted through 12 hours of RBCC:Nb^mCh^ expression (mean fluorescence intensity 25±18%) (Figure 5H and I). Hht2:mCh degradation was also measured by immunoblotting. Hht2:mCh protein levels were 108±62% and diminished to 70±5% at one and six hours after RBCC:Nb^mCh^ expression, respectively (Supplemental Figure 4C and D). Thus, *Tet* Trim-Away can degrade nuclear proteins but degradation can take up to three hours.

### *Tet* Trim-Away elicits protein depletion phenotypes

Rapid depletion of specific proteins by *Tet* Trim-Away would be expected to result in phenotypes related to those previously detected in mutant cells with complete knockouts of the corresponding genes. To test this idea, we analyzed the phenotypes of cells depleted for proteins previously shown by gene knockout studies to be non-essential (Poc1) and essential (Bld10) (Bayless et al., 2012; Pearson et al., 2009). For these studies, the transgenes encoding Poc1:mCh and Bld10:mCh fusions were driven to complete fixation by phenotypic assortment, completely replacing the wildtype genes (Merriam and Bruns, 1988; Ruehle et al., 2016). Thus, the cells contained no pool of the untagged (and therefore depletion-resistant) protein of interest.

RBCC:Nb^mCh^ was expressed in Poc1:mCh cells at both 30°C, the normal growth condition for this species, but also 39°C. The higher temperature was tested because cells with complete knockout of the *POC1* gene (*poc1Δ*) were previously shown to be thermosensitive, exhibiting severe morphological and motility defects at 39°C (Pearson et al., 2009). After six hours of RBCC:Nb^mCh^ induction at 39°C, 92±9% cells exhibited no detectable Poc1:mCh signal (mean fluorescence intensity reduced to 25±13% of control cells; Figure 6A-C). There was also visible loss of centrin foci, a proxy for basal bodies (Figure 6A and D) which became even more pronounced at 12 and 24 hours. In cultures induced for 24 hours, cells were no longer dividing (Figure 6A-D) and accumulated centrin-negative autofluorescent puncta in the cytoplasm that precluded measurement of the extent of protein depletion at this timepoint. Control cells at 39°C, i.e., not exposed to CdCl_2_, exhibited mild defects in basal bodies and cortical row structure, possibly due to leaky expression of the RBCC:Nb^mCh^ degron (Figure 6A and C).

**Figure 6.**
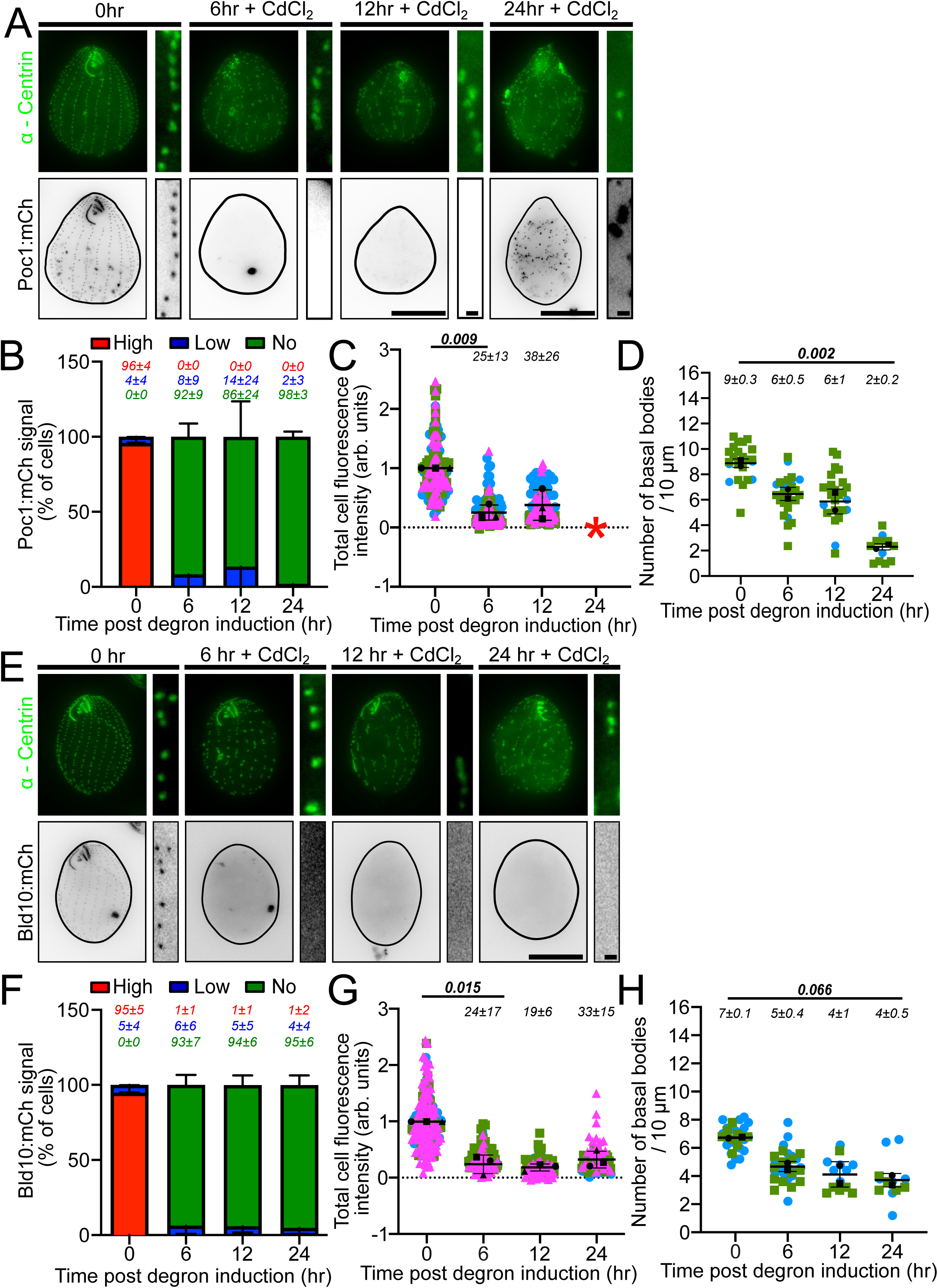
*Tet* Trim-Away elicits mutant phenotypes. (A) Cells at 39°C stained with α-Centrin (green) to visualize basal bodies on top and Poc1:mCh in grey scale (equally scaled). Centrin images are not equally scaled. For representative images of cells at 12 hours, the contrast is increased by 42% and the brightness is decreased by 26% as compared to images of cells at 0, 6, and 24 hours. For representative images of insets of cells at 12 hours, the contrast is increased by 44% and the brightness is decreased by 41% compared to insets of cells images of cells at 0, 6, and 24 hours. (B) Qualitative analysis of Poc1:mCh degradation. (C) Quantitative analysis of full cell Poc1:mCh mean fluorescence intensity through 24 hours of RBCC:Nb^mCh^ Trim-Away expression. (D) Mean number of basal bodies per 10 μm through 24 hours of RBCC:Nb^mCh^ Trim-Away expression. Scale bar for images of 0, six, and 12 hour images is 20μm and 1μm and scale bar for images of cells at 24 hours is 30μm and 1μm. (E) Representative images of Bld10:mCh cells expressing RBCC:Nb^mCh^ Trim-Away. α-Centrin staining to visualize basal bodies (green) are not equally scaled. For representative images of cells at 0 and 12 hours, the contrast is increased by 56% and the brightness is decreased by 35% as compared to images of cells at other time points. Bld10:mCh cells in grey scale (bottom, equally scaled). (F) Qualitative analysis of Bld10:mCh degradation through 24 hours of RBCC:Nb^mCh^ induction. (G) Whole cell Bld10:mCh mean fluorescence intensity through 24 hours of RBCC:Nb^mCh^ Trim-Away induction. (H) Mean number of basal bodies per 10 μm through 24 hours of RBCC:Nb^mCh^ Trim-Away expression. Percentages of each depletion class and mean fluorescence intensity of each time point relative to control are indicated above the data points. Scale bar, 20μm, 1μm. Error bars indicate mean±SD. (B, C, F, G) *N > 300* for qualitative analysis and *N > 120* for quantitative analysis. (D, H) *N = 5 basal body rows / cell and N > 12 cells*.

Similar to the previously characterized *poc1Δ* knockout cells, the phenotypes in the induced RBCC:Nb^mCh^ cells were temperature sensitive. That is, when RBCC:Nb^mCh^ expression was induced in Poc1:mCh cells at 30°C rather than 39°C, Poc1:mCh signal was depleted within six hours but without accompanying loss of basal bodies (Supplemental Figure 5A-D). Mild basal body disorganization was observed at six, 12 and 24 hours, but without defects in cell division or cell survival. Taken together, our results indicate that *Tet* Trim-Away depletion of Poc1 rapidly phenocopies both the defects of *poc1Δ* cells and their thermosensitivity.

Bld10 is an essential basal body protein since *bld10Δ* gene knockout mutants fail to divide. We tested whether *Tet* Trim-Away depletion of Bld10 could similarly reveal its essentiality. After six hours of RBCC:Nb^mCh^ induction in Bld10:mCh cells, 94±7% of cells had no detectable Bld10:mCh. The mean fluorescence intensity was reduced to 25±13% of control cells throughout the time course (Figure 6E-H). Importantly, Bld10 Trim-Away cells lack basal bodies and stop dividing by 24 hours (Figure 6E and H) and thus phenocopy the genetic knockout of Bld10. Cells at six hours phenocopied the 12 hour timepoint of *bld10Δ* genetic knockout (Bayless et al., 2012). Therefore, degradation of homozygous targeted proteins by induction of *Tet* Trim-Away leads to the rapid detection of phenotypes consistent with those observed in genetic knockouts.

## DISCUSSION

*Tet* Trim-Away is a rapid and selective protein degradation system in *Tetrahymena* that destroys proteins within minutes, maintains degradation for weeks, and is reversible. RBCC:Nb^mCh^ localizes to the site of the target protein and does not cause detectable collateral damage. *Tet* Trim-Away can be used to target cytoplasmic, basal body, ciliary, and nuclear proteins, and it renders loss-of-function phenotypes in essential and non-essential genes.

Functional studies of specific proteins in *Tetrahymena* have largely relied on gene knockouts. In one important approach, the gene of interest is knocked out in the germline (silent) micronucleus while the somatic (expressed) macronucleus retains the intact gene (KO heterokaryons) (Hai and Gorovsky, 1997). The phenotype is not realized until heterokaryons are mated and drug resistance selects for progeny bearing the now-homozygous knockout allele in the expressed macronucleus. This approach requires significant time for both strain construction and genetic manipulations, and multiple cell cycles before phenotypic analysis is possible, during which compensatory pathways can arise and obscure the direct consequences of the protein loss. In comparison, *Tet* Trim-Away has three major advantages. Most importantly, extensive depletion of the targeted protein is rapid, occurring within an hour for some examples tested in this paper. *Tet* Trim-Away degradation may be more efficient in producing loss-of-function phenotypes than genomic knockouts (Figure 6E-G), perhaps due to expansion of compensatory mechanisms in the latter. Secondly, the level of protein degradation can be readily measured as changes in mCh fluorescence or immunoreactivity and correlated with any defects that are noted. One example of the usefulness of this comes from our results with *Tet* Trim-Away-mediated depletion of Bld10:mCh, where the phenotypes we observed after six hours of degron induction matched those of *bld10Δ* cells at 12 hours post genomic knockout. Third, construction of the *Tet* Trim-Away strains only involves macronuclear transfection with an mCh-tagged allele for a protein of interest, and stable integration of the RBCC:Nb^mCh^ cassette, both of which are straight-forward in *T. thermophila*. As long as the tagged allele is functional, it can be driven to fixation at the endogenous macronuclear locus just by applying increasing selective pressure via a linked drug-resistance cassette.

A limitation to *Tet* Trim-Away is that some target proteins in our studies were incompletely depleted. One possible explanation is that cells in different stages of the cell cycle maintain different mechanisms of protein degradation. For example, starvation-arrested cells in G1 do not effectively degrade proteins targeted with *Tet* Trim-Away (data not shown), raising the possibility that one or more steps in proteosome-dependent degradation is downregulated in G1. Consistent with this, genes annotated as proteasome function are reduced during G1 and S-phase (Bertagna et al., 2025). An additional consideration is that a degron may promote elevated levels of the ubiquitinylated target protein, reaching concentrations that ultimately exceeds the proteasomal capacity. Here, abundant proteins in the cell may saturate the proteasome thereby limiting both access to the *Tet* Trim-Away cargo as well as normal cellular functions of the proteasome. This has been observed with proteasomal overload in photoreceptor and neuronal cells (Cenci et al., 2006; Dantuma and Lindsten, 2010; Lindsten and Dantuma, 2003; Lobanova et al., 2013).

*Tet* Trim-Away function relies on interactions with *Tetrahymena* E1’s and E2’s, to ubiquitinylate the target protein. No TRIM21 homolog can be detected in the *Tetrahymena* genome, and we presume that *Tetrahymena* E1 and E2 enzymes work with the exogenous RBCC. In vertebrates, several E1 and E2 enzymes like UBE2D1/UbcH5a, UBE2E1/UbcH6, UBE2W, UBE2N, UBE2D2 UbcH5b, UBE2D3/UbcH5c interact with human TRIM21 (Anandapadamanaban et al., 2019; Fletcher et al., 2015a; Fletcher et al., 2015b; Vittal et al., 2015). Importantly, the *Tetrahymena* genome has predicted orthologs of some of these E1 and E2 enzymes (Eisen et al., 2006). Further studies are required to understand which *Tetrahymena* E1 and E2 enzymes promote *Tet* Trim-Away activity and whether these could be limiting factors for *Tet* Trim-Away-mediated degradation.

Our studies suggest that there is a low level of RBCC:Nb^mCh^ expression even in uninduced cultures (i.e., no CdCl_2_) (Figure 3C) and subsequent target protein degradation and turnover (Supplemental Figure 4C). Consistent with this, mild mutant phenotypes were observed in the homozygous Poc1:mCh *Tet* Trim-Away strains under non-induced conditions (Figure 6A). Variants on the *Tet* Trim-Away construct we have introduced here, or similar efforts, may benefit from tighter protein expression control and/or the use of alternative mechanisms for recognizing the protein of interest. One such strategy may be to express rapamycin-inducible fusions of RBCC:FKBP with a target protein fused to FRB and use both transient RBCC:FKBP expression with rapamycin for selective expression and protein degradation.

## RESOURCE AVAILABILITY

Reagents used in this project will be available upon request.

## AUTHOR CONTRIBUTIONS

Conceptualization, GD, AJSW, AA, APT, CGP; Investigation, GD, AJSW, AA, CGP; Analysis, GD, AA; Writing, GD, CGP; Reviewing and Editing, GD, AJSW, APT, CGP; Funding, CGP.

## ACKNOWLEDGEMENTS

The authors are thankful to all members of the Pearson Lab for their support and guidance. CGP is funded by NIGMS R35 GM140813 and NSF 2421873.

## MATERIALS AND METHODS

### *Tetrahymena thermophila* cell lines and culturing

Strains used in this study are IA267, B1868, SB1969 which were obtained from The National Tetrahymena Stock Center (Washington University, formerly Cornell University) and Rescue 1:Poc1:mCh (Stemm-Wolf et al., 2026). Cells were grown at 30°C in 2% SPP (2% proteose peptone, 0.2% glucose, 0.1% yeast extract and 0.003% Fe-EDTA in HPLC-grade water (Thermo Fisher, Waltham, MA, USA) unless otherwise indicated. Cells were counted with a Z1 Coulter Counter (Beckman Coulter, Brea, CA). Experiments were conducted with mid log phase cultures (100,000 – 500,000 cells/mL).

### Plasmids

All the mCh and GFP fusion proteins used in this study are described in Table 1. Constructs were generated by cloning ∼1 kilobase flanking regions at the C-terminal ends of the coding sequences for homologous recombination at the endogenous loci. Drug selections for insertions are described in Table 1. Drug stock concentrations for selection are – Paromomycin = 200 μg/mL and Cycloheximide = 15 μg/mL. Cells were typically selected for one drug at a time, with paromomycin first followed by cycloheximide.

The pBS-MTT1-RBCC:Nb^mCh^ or pBS-MTT1-RBCC:Nb^GFP^ constructs were designed using the human RBCC sequence with a FLAG tag at the C-terminal end. Nanobody sequences specific for mCh or GFP (Fridy et al., 2014; Holzer et al., 2022) were attached at the C-terminal end of the RBCC motif. This plasmid integrates at the *RPL29* locus and confers cycloheximide (cyhx) resistance. All RBCC:Nb strains were assorted to and maintained at 30 μg/mL cyhx (Bleyman and Bruns, 1977). Expression of *Tet* Trim-Away was induced using the *MTT1* promoter with 5.5 μM (1.0 μg/mL) CdCl_2_ (Shang et al., 2002).

### Macronuclear transformation

Strains were generated by transforming linearized constructs via biolistic bombardment into the macronucleus. 50 mL of cells were grown in SPP to ∼200,000 cells/mL and starved in 10mM Tris, pH 7.4 overnight (Orias et al., 1999). A PDS1000 biolistic particle system with 900 psi rupture disks and 0.6 μm Tungsten particles (Bio-Rad, Hercules, CA) was used for transformations (Bruns and Cassidy-Hanley, 2000).

### Immunofluorescence

For image analysis, 750,000 – 1,500,000 cells were washed with 10 mM Tris, pH 7.4 and fixed for 5-20 minutes in PHEM (PIPES, HEPES, EGTA, MgCl_2_) with 3.2% Paraformaldehyde and 0.24% TritonX-100, followed by a 10 minute permeabilization in X% TritonX-100 in PHEM on ice. (Canman et al., 2000). Cells were washed 3X with 0.5% BSA in PHEM and stored at 4°C in 0.5% BSA in PHEM until imaging. For immunofluorescence staining, cells were stained with 1:200 α-Flag ANTI-FLAG M2 antibody (mouse monoclonal, Sigma-Aldrich, Merck Group, St. Louis MO, USA),1:1000 α-CEN1 2111 (Stemm-Wolf et al., 2005) or 1:800 6-11-B1 α-acetylated α-tubulin (Blasius et al., 2025). Secondary antibodies used were 1:1000 anti-Mouse Alexa 488, 1:1000 anti-Rabbit Alexa 488, 1:1000 anti-Rabbit Alexa 647 (Thermo Fisher, Waltham, MA, USA and Jackson ImmunoResearch Labs, West Grove, PA, USA). Primary and secondary antibodies were diluted in freshly made 0.5% BSA in PHEM and the incubations were carried out at room temperature for 1.5 hours or overnight at 4°C and 1 hour at room temperature or 2 hours at 4°C, for primary and secondary antibody incubations, respectively. Cells were washed 3X with 0.5% BSA in PHEM after each antibody incubation. 6.5 μl of the final pellet of cells was mixed with 5 μl antifade (1X PBS, 90% Glycerol, 0.5% n-propyl Gallate) and put on slides with a glass cover slip and sealed using clear nail polish.

B1868 or B2086 p4T2-1-Poc1-GFP-NEO2 cells were fixed using 70% ethanol with 0.12% TritonX-100. Cells were placed on ice for five minutes followed by washing with 10mM Tris, pH 7.4 and addition of 5 ml of ice-cold fixative. Cells were kept on ice for 20 minutes after which they were washed 3X with 5 mls of 0.5% BSA in TBS and stored at 4°C for imaging.

### Imaging

Cells were imaged at room temperature on a Nikon Eclipse Ti inverted microscope using a 100X Plan Apochromat oil objective (NA 1.45) (Nikon Instruments, Inc., Melville, NY, USA) and an Andor iXon X3 camera (Oxford Instruments, Abingdon, UK). Cells were also imaged using a Nikon Eclipse Ti inverted microscope stand equipped with a 100x Plan Apochromat oil objective (NA 1.45) (Nikon Instruments, Inc., Melville, NY, USA), a Teledyne Photometrics Prime95B sCMOS camera (Teledyne Photometrics), and CSU-X1 (Yokogawa Electric Corporation, Sugar Land, TX, USA) spinning disk. Slidebook 2025 digital microscopy software (Intelligent Imaging Innovations, Inc, Denver, CO, USA) was used for image acquisitions.

### Image analysis

To quantify the signal intensity of the fluorescently labelled proteins within cells, regions of interest (ROIs) of cells or basal bodies were generated followed by ROIs of three background regions. Fluorescence intensity analysis of cells or basal bodies was performed using integrated density (ID) values in ImageJ of maximum projected z-stacks images. To account for variability, the three background ROIs ID values were averaged and subtracted from the ID of the cell, after adjusting for the difference in size of the ROIs. The ID was then normalized for each cell or basal body to the average ID value of the control cells or basal bodies in the given experiment.

### Statistical Methods

Data was organized in and analyzed using Microsoft Excel, R-studio (Posit, Boston, MA, USA), and Graphpad Prism (GraphPad Software, Boston, MA, USA). All experiments had at least two biological replicates and the normalized values of each cell or basal body from each replicate is differentiated based on the data points shape and color in the graphs. Error bars represent the standard deviation and data points in black represent normalized averages of each biological replicate. The Student’s independent unpaired t-test (performed in R-studio) was used to determine the significance between two normal unpaired samples. Differences between distributions were considered statistically significant if the *p* values were less than 0.05.

### Western blots

1,000,000 – 1,500,000 cells were lysed in buffer (50 mM Tris-HCl, 150 mM Sodium Chloride, 20 mM EDTA, 1% TritonX-100, 10 mM β-mercaptoethanol, 1% Sodium deoxycholate) with 5 mM PMSF, and 1X cOmplete Protease Inhibitor Cocktail (Sigma-Aldrich, Merck Group, St. Louis MO, USA). . Approximately 45 μg of protein was used. Western blots were blocked with TBS + 0.05% Tween-20 (TBST) and 5% non-fat milk for one hour at room temperature and incubated with primary antibody (1:1000, Living colors mouse monoclonal antibody, Takara Bio USA, San Jose, CA and 1:5000, Anti-beta Tubulin antibody- Loading Control, Abcam, Danaher Corporation, Cambridge, UK) for two hours at room temperature. Blots were incubated in Licor IR680 and IR800 secondary antibodies at 1:10,000 for one hour. Blots were washed three times with TBST after each antibody incubation. Blots were imaged using fluorescence Odyssey CLx Imager and LI-COR Acquisition Software (LI-COR Biotechnology Lincoln, NA, USA).

## SUPPLEMENTAL FIGURE LEGENDS

**Supplemental Figure 1. Tet Trim-Away for Poc1:mCh**

(A) A schematic of the *Tet* Trim-Away construct. The Ring, B-Box, and Coiled Coil domains of TRIM21 are attached to a nanobody specific to mCh or GFP. This construct is driven by the metallothionein (MTT1) promoter which is responsive to CdCl_2_. The flag tag is on the C-terminal end of the RBCC motif. (B) CdCl_2_ dosage analysis of RBCC:Nb^mCh^ Trim-Away in Poc1:mCh cells. Cells were exposed to 0.1, 0.5 and 1.0 μg/ml CdCl_2_ for up to six hours and analyzed for Poc1:mCh depletion. (C) Whole cell Poc1:mCh fluorescence intensity for cells exposed to 0.1, 0.5 and 1.0 μg/ml CdCl_2_ for up to six hours. Percentages of each depletion class and mean fluorescence intensity of each time point relative to control are indicated above the data points. (D) Wildtype cells with RBCC:NB^mCh^ Trim-Away stained with α-centrin in green. Scale bar, 20μm. Error bars indicate mean±SD. (B,C) *N > 400* for qualitative analysis and *N > 160* for quantitative analysis.

**Supplemental Figure 2. Prolonged Tet Trim-Away degradation of Poc1:mCh and degradation of Poc1:GFP by RBCC:NbGFP**

(A) Qualitative analysis of RBCC:Nb^mCh^ Trim-Away degradation of Poc1:mCh over 336 hours. (B) Whole cell Poc1:mCh mean fluorescence intensity of RBCC:Nb^mCh^ Trim-Away degradation of Poc1:mCh through 336 hours of induction. (C) Qualitative analysis of RBCC:Nb^GFP^ Trim-Away degradation of Poc1:GFP through 12 hours of induction. (D) Whole cell Poc1:GFP mean fluorescence intensity of cells through 12 hours RBCC:Nb^GFP^ induction. Percentages of each depletion class and mean fluorescence intensity of each time point relative to control are indicated above the data points. Error bars indicate mean±SD. (A, B, C, D) *N > 300* for qualitative analysis and *N > 120* for quantitative analysis.

**Supplemental Figure 3. Tet Trim-Away localizes to basal bodies**

(A). α-Flag immunofluorescence staining (green) in wild type cells that lack RBCC:Nb^mCh^ Trim-Away and lack Poc1:mCh (channel shown in grey scale). (B) Wildtype cells with RBCC:NB^mCh^ Trim-Away but lacking Poc1:mCh (channel shown in grey scale) stained with α-Flag (green). (C) Poc1:mCh cells (grey scale) without RBCC:Nb^mCh^ Trim-Away stained with α-Flag (green). MG132 was added to cells 45 minutes before addition of CdCl_2_. Cells were fixed and imaged one hour after CdCl_2_ treatment. Scale Bar, 20μm.

**Supplemental Figure 4. Tet Trim-Away degrades proteins in diverse cellular compartments**

(A) Western blot of Bbc39:mCh degradation at 0, one and six hours of RBCC:Nb^mCh^ induction. Bbc39:mCh in green and loading control in red. (B) Western blot of Bbc39:mCh degradation through six hours of RBCC:Nb^mCh^ induction and quantification of Bbc39:mCh protein degradation. (C) Western blot of Hht2:mCh degradation at 0, one and six hours of RBCC:Nb^mCh^ induction. Hht2:mCh in green and loading control in red. Positive control is Hht2:mCh cells without RBCC:Nb^mCh^ Trim-Away (marked as PC). (D) Western blot of Hht2:mCh degradation through six hours of RBCC:Nb^mCh^ induction and quantification of Hht2:mCh protein degradation. Error bars indicate mean±SD.

**Supplemental Figure 5. Tet Trim-Away elicits mutant phenotypes**

(**A**) Homozygous Poc1:mCh cells at 30°C stained with α-Centrin (green) to visualize basal bodies on top and Poc1:mCh in grey scale at the bottom (all images are equally scaled).

(**B**) Qualitative analysis of Poc1:mCh degradation. (C) Quantitative analysis of full cell Poc1:mCh mean fluorescence intensity through 24 hours of RBCC:Nb^mCh^ Trim-Away expression. Percentages of each depletion class and mean fluorescence intensity of each time point relative to control are indicated above the data points. (D) Mean number of basal bodies per 10 μm through 24 hours of RBCC:Nb^mCh^ Trim-Away expression. Scale bars, 20μm and 1μm. Error bars indicate mean±SD. (B, C) *N > 300* for qualitative analysis and *N > 120* for quantitative analysis. (D) *N = 5 basal body rows / cell and N > 12 cells*.

## Abbreviations

RBCC:Nb^mCh^: Ring-B-box-coiled coil domain with mCh nanobody
CdCl_2_ cadmium chloride: metallothionein (MTT1)

