## Supplemental Figures for "*Tet* Trim-Away: A conditional rapid protein degradation system for *Tetrahymena thermophila*"

### Supplemental Figure 1

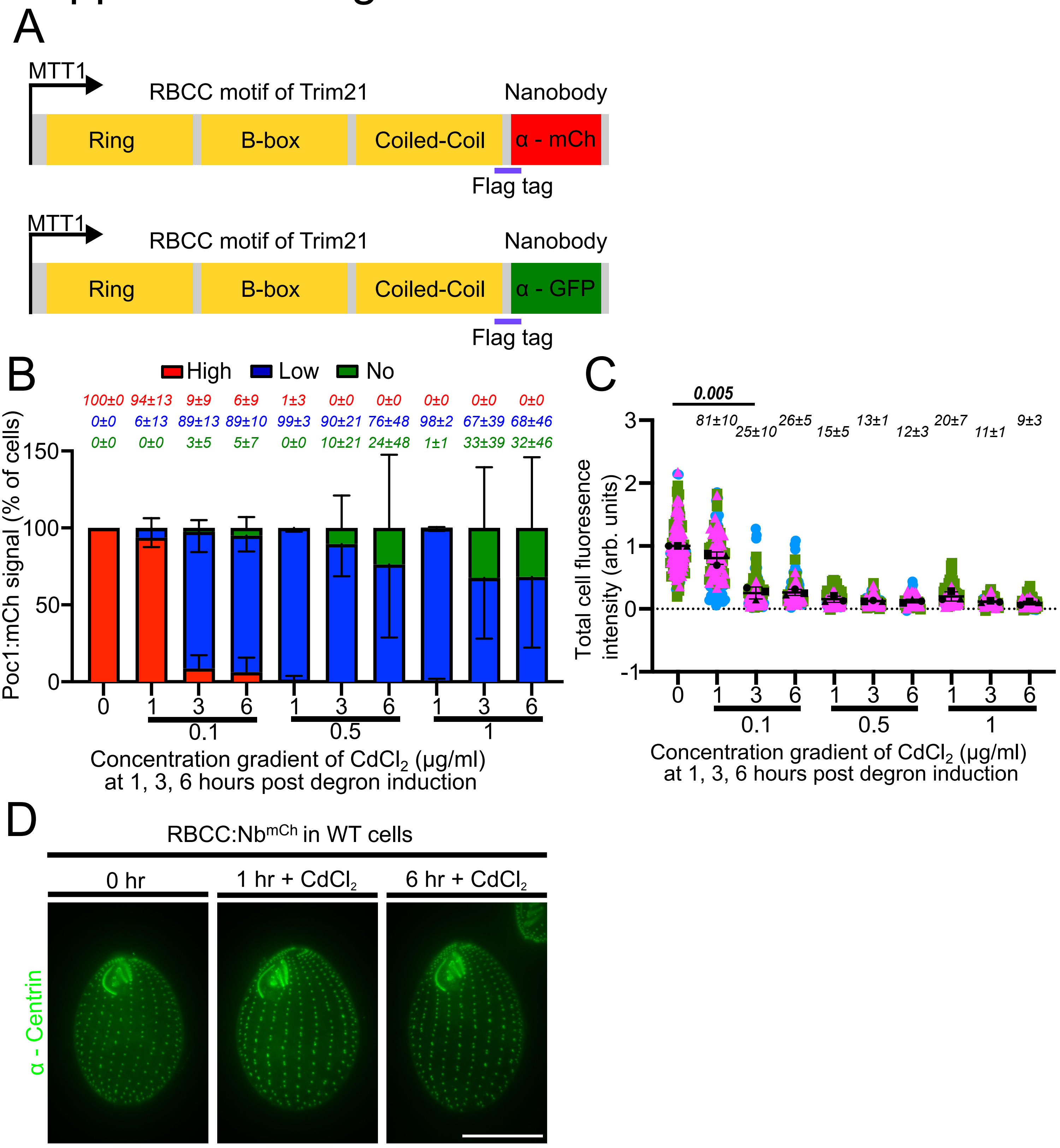

### Supplemental Figure 2

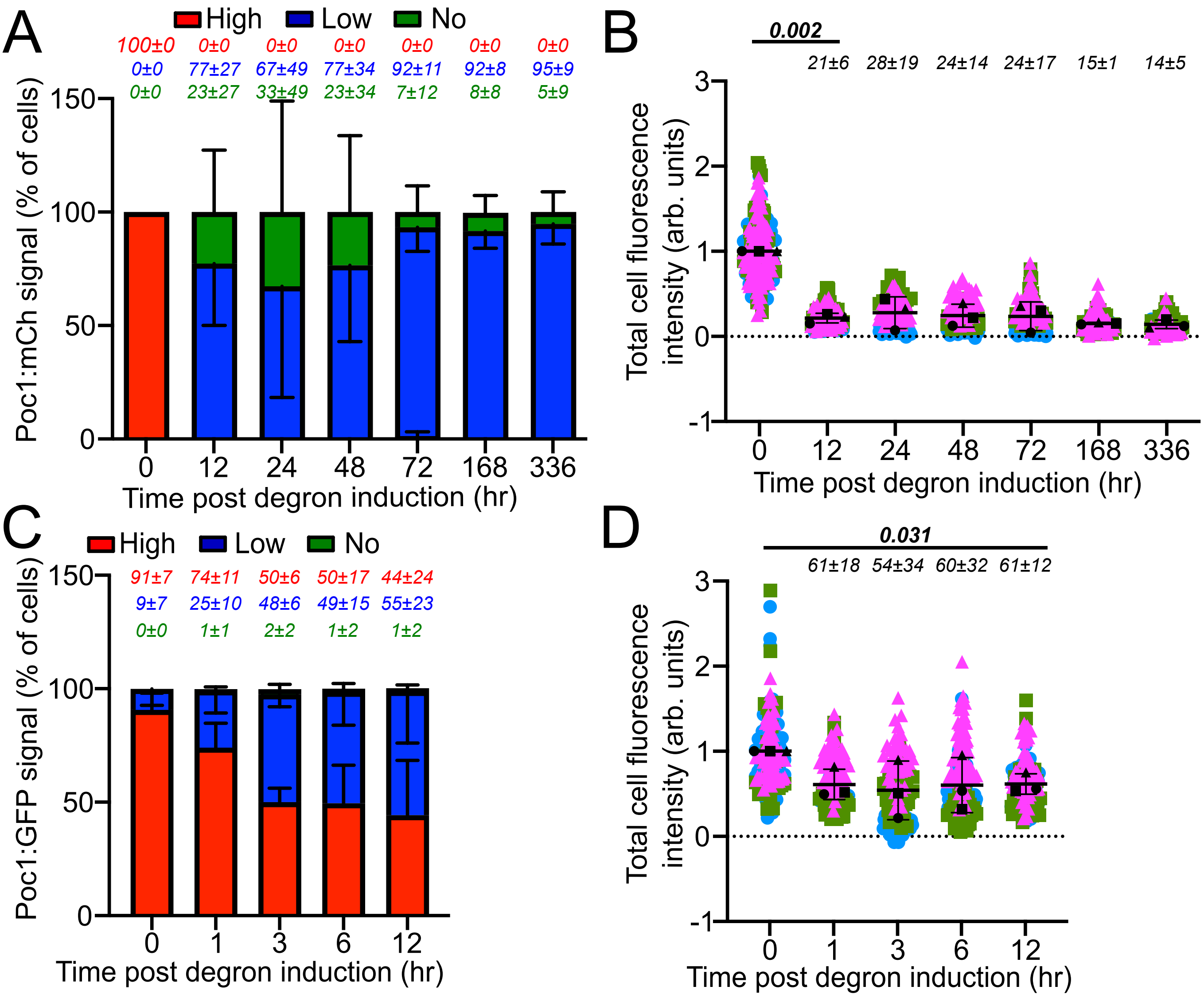

### Supplemental Figure 3

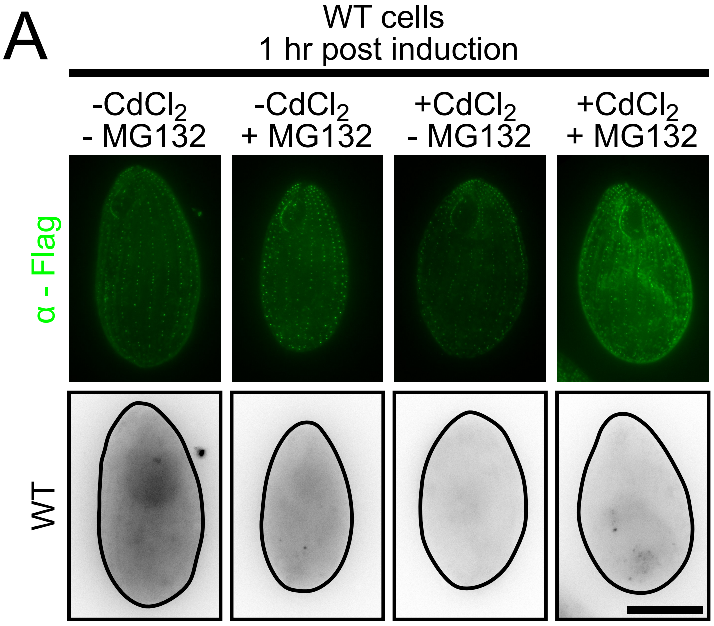


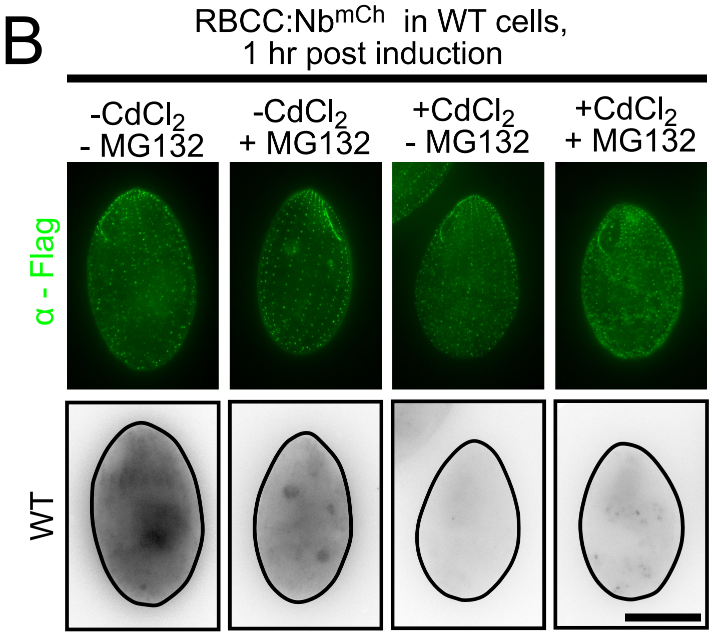


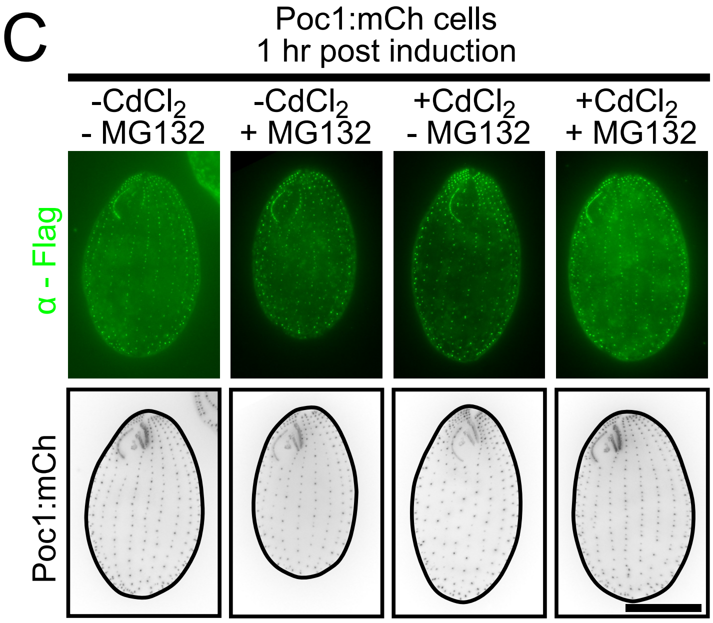

### Supplemental Figure 4

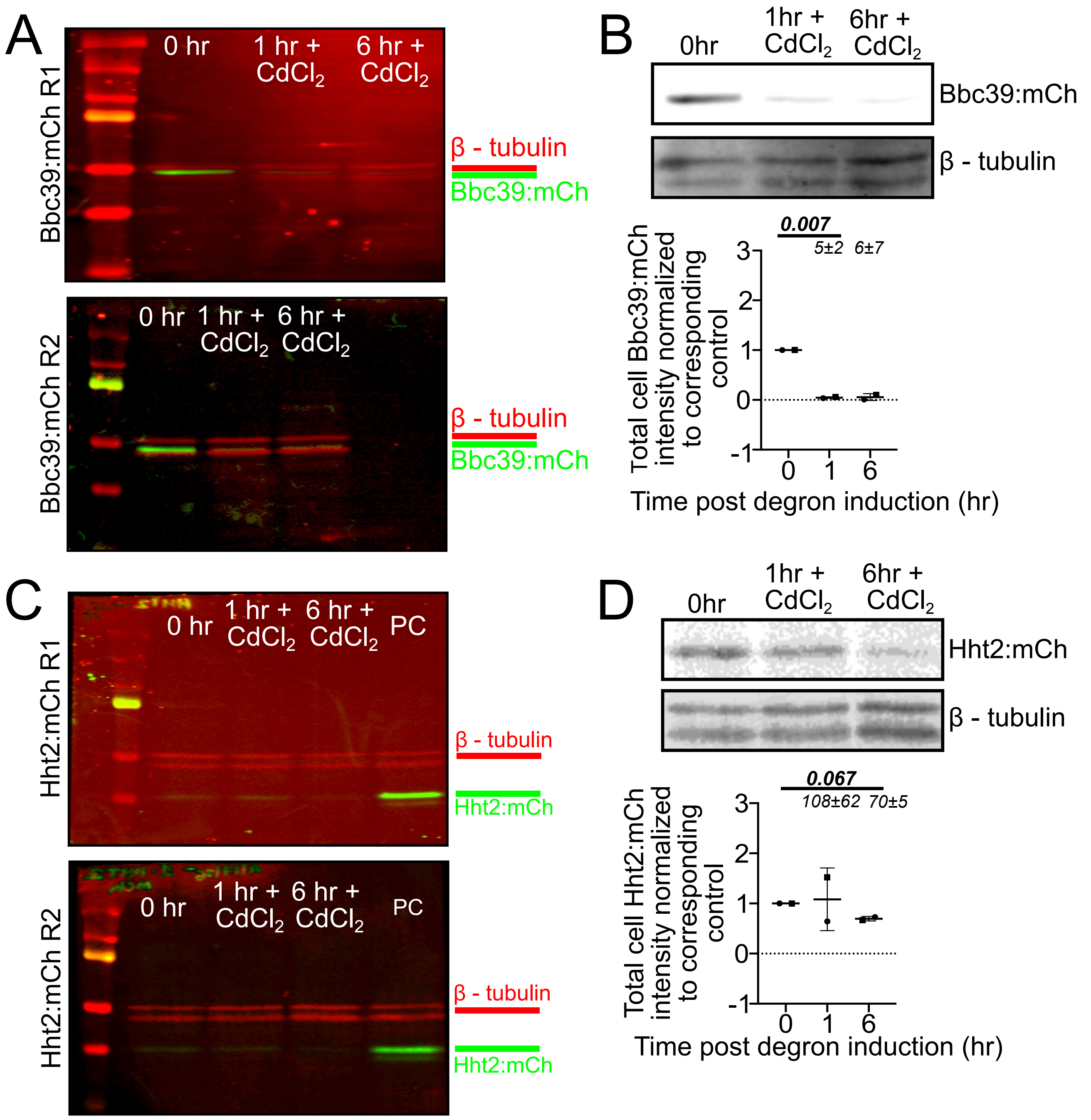

### Supplemental Figure 5

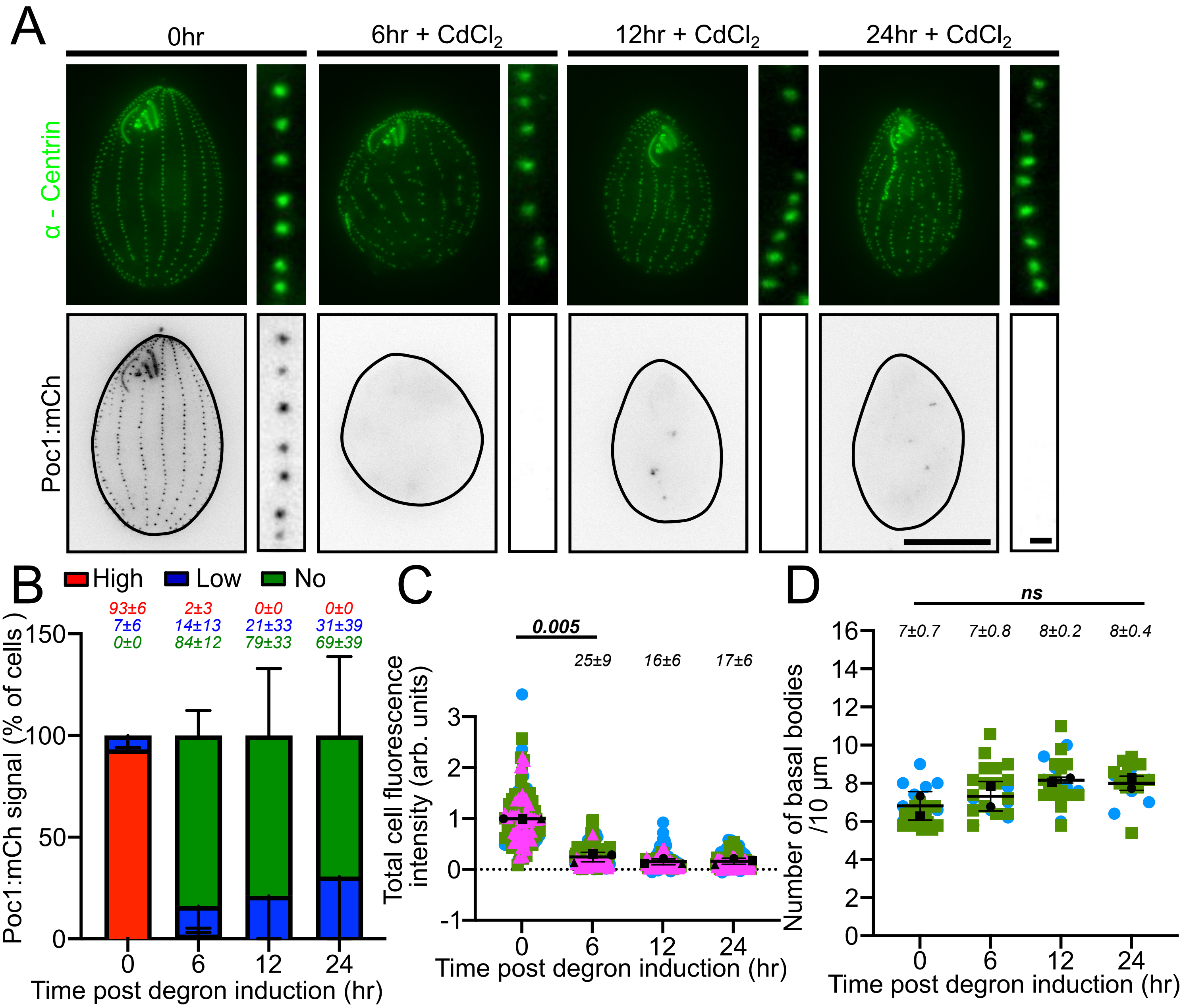
